# Graph Contrastive Learning as a Versatile Foundation for Advanced scRNA-seq Data Analysis

**DOI:** 10.1101/2024.04.23.590693

**Authors:** Zhenhao Zhang, Yuxi Liu, Meichen Xiao, Kun Wang, Yu Huang, Jiang Bian, Ruolin Yang, Fuyi Li

**Author notes:** Zhenhao and Yuxi are equal contributors to this work.

## Abstract

Single-cell RNA sequencing (scRNA-seq) offers unprecedented insights into transcriptome-wide gene expression at the single-cell level. Cell clustering has been long established in the analysis of scRNA-seq data to identify the groups of cells with similar expression profiles. However, cell clustering is technically challenging, as raw scRNA-seq data have various analytical issues, including high dimensionality and dropout values. Existing research has developed deep learning models, such as graph machine learning models and contrastive learning-based models, for cell clustering using scRNA-seq data and has summarized the unsupervised learning of cell clustering into a human-interpretable format. While advances in cell clustering have been profound, we are no closer to finding a simple yet effective framework for learning high-quality representations necessary for robust clustering. In this study, we propose scSimGCL, a novel framework based on the graph contrastive learning paradigm for self-supervised pretraining of graph neural networks. This framework facilitates the generation of high-quality representations crucial for cell clustering. Our scSimGCL incorporates cell-cell graph structure and contrastive learning to enhance the performance of cell clustering. Extensive experimental results on simulated and real scRNA-seq datasets suggest the superiority of the proposed scSimGCL. Moreover, clustering assignment analysis confirms the general applicability of scSimGCL, including state-of-the-art clustering algorithms. Further, ablation study and hyperparameter analysis suggest the efficacy of our network architecture with the robustness of decisions in the self-supervised learning setting. The proposed scSimGCL can serve as a robust framework for practitioners developing tools for cell clustering. The source code of scSimGCL is publicly available at https://github.com/zhangzh1328/scSimGCL.

## Introduction

Cells are essential building blocks of all living organisms and play a pivotal role in a myriad of biological functions. The heterogeneity in gene expression across individual cells has emerged as a significant field of study, especially with the development of single-cell RNA sequencing (scRNA-seq). The scRNA-seq, perhaps the most widely used method for transcriptome-wide analysis at the single-cell level [1], allows a deeper insight into cell classification, characterization, and differentiation. Such analysis have successfully uncovered the complex association between cellular functions and improved the performance of various downstream applications. These applications include profiling genetic diversity within populations [59], identifying and annotating cell subtypes [52], and facilitating advances in drug discovery and development [38].

However, raw scRNA-seq data present their own set of challenges, such as high dimensionality and measurement noise [31]. Notably, measurement noise often causes dropout events, where genes appear to be unexpressed due to technical artifacts rather than biological reality. Recent advancements in deep learning techniques have shown promise in addressing these challenges by reducing the dimensionality of data and filtering out the noise to uncover biological signals. Deep learning techniques have been widely used in bioinformatics, including scRNA-seq data analysis, to provide new insights into cellular functions.

In the realm of scRNA-seq data analysis, deep neural networks have been usefully explored. Prior studies by [35, 23, 9, 53, 36, 15, 14, 25] have employed autoencoders – a type of neural network to effectively learn condensed representations of scRNA-seq data in a lower-dimensional space. Representative examples of scDeepCluster, DESC, and scDCC [35, 23, 36], which are specifically designed to provide clustering assignments with scRNA-seq data. Such approaches, DCA and scGMAI [9, 53], integrate data imputation into their clustering assignments to mitigate the impact of dropout events, thereby facilitating the formation of informative gene expression matrices, which is essential for improving cell clustering.

More recent studies have seen increasingly rapid advances in the field of scRNA-seq data analysis [7, 44, 4, 6, 42, 10]. On the one hand, graph-sc, scASGC, and scGAC [7, 44, 4] employ graph autoencoders to encode scRNA-seq data as cell graphs and capture the between-cell association. On the other hand, contrastive-sc, scDCCA, and scDECL [6, 42, 10] focus on the provision of autoencoder with contrastive learning. The intuition behind contrastive learning is to compare input samples in order to learn desirable representations using the similarity between positive pairs or the dissimilarity between negative pairs. Accordingly, contrastive learning is recognized as an alternative approach to reconstruction-based with a focus on learning an embedding space where similar ones are pushed closer to each other while dissimilar ones are pushed farther apart [11, 12, 27, 56].

Despite these promising advances, research has yet to systematically develop a simple yet effective framework for learning high-quality representations crucial for robust cell clustering. The learning of high-quality representations is built upon the success of pretraining generalizable models, which aligns with the promise of the current foundation models (i.e., a series of large-scale pretrained models that can be applied on various downstream use cases and tasks) [29, 47, 30]. The foundation model has been instrumental in our understanding of the role of deep learning in the biological context. However, the transition to practical, actionable insights in scRNA-seq remains challenging, especially in analyzing and interpreting complex biological data.

To bridge this gap, we incorporate the graph contrastive learning (GCL) paradigm into scRNA-seq data analysis. Several key aspects of modeling need to be carefully considered. First, the construction of cell-cell graphs from scRNA-seq data presents a fundamental challenge. Second, there is an urgent need to design a neural network architecture that leverages the GCL paradigm to learn representations desirable for cell-cell graphs. Third, the development of pretraining strategies as a pretext task in the self-supervised learning setting is critical for improving the effectiveness and robustness of the model decisions.

In this study, we propose scSimGCL, a simple and effective framework that combines graph neural networks with contrastive learning, aligning with the GCL paradigm, specifically tailored for scRNA-seq data analysis. The GCL paradigm facilitates self-supervised pretraining of graph neural networks, which enables the generation of high-quality representations crucial for robust cell clustering. A critical component of our scSimGCL is the innovative construction of cell-cell graphs using scRNA-seq data. We develop a cell-cell graph structure learning mechanism that pays attention to the critical parts of the input data using a multi-head attention module [40] for improving the accuracy and relevance of graphs. This module is used for the purpose of deriving a nuanced cell-cell graph, where individual cells form nodes and their biological associations are represented as edges.

Moreover, our study addresses a major issue in existing methods that combine graph machine learning with contrastive learning for scRNA-seq data. Prior studies [19, 34, 50] often directly integrate preprocessed cell graphs (e.g., via K-Means) into the contrastive learning scheme. However, this method is likely to disrupt the intrinsic homophily of the graph, where connected nodes are expected to share similar properties, thus resulting in inferior quality of representation. In response to this issue, our approach adopts an effective strategy for constructing the pairs of contrastive learning using the homophily of original and augmented cell-cell graphs (generated by gene masking and edge dropping [26]). This strategy preserves the internal structure of graphs, facilitating the generation of informative contrastive pairs. Further, our approach introduces a well-designed pretraining mechanism to explore the topological nuances of cell-cell graphs. This mechanism implements contrastive learning with data imputation in a joint learning strategy, enhancing the model performance across various evaluation metrics.

We conduct rigorous evaluations of scSimGCL against deep learning approaches on both simulated and real scRNA-seq datasets. Our framework outperforms competing state-of-the-art models in scRNA-seq data imputation and cell clustering by a considerable margin. Further experimental investigations into the clustering assignments reveal that scSimGCL can seamlessly integrate state-of-the-art clustering algorithms into its architecture. Notably, the model is capable of making informed decisions by indicating a predefined number of clusters or automatically determining the number of clusters (i.e., K-Means and Affinity Propagation), which is a key aspect in unsupervised learning scenarios such as scRNA-seq data analysis. Ablation study and hyperparameter analysis further confirm the efficacy and robustness of scSimGCL in the self-supervised learning setting.

## Materials and methods

### The framework of scSimGCL

The overall structure of the proposed scSimGCL is illustrated in Figure 1. First, we incorporate a multi-head attention module [40] to learn a cell-cell graph from scRNA-seq data. Second, we adopt feature/gene masking and edge dropping in the learned cell-cell graph to generate its counterpart. Subsequently, the learned cell-cell graph and its counterpart are used to create contrastive pairs. Last, we design a pretraining mechanism to generate high-quality representations for cell clustering, where contrastive learning and data imputation are implemented with a joint learning strategy.

**Fig. 1.**
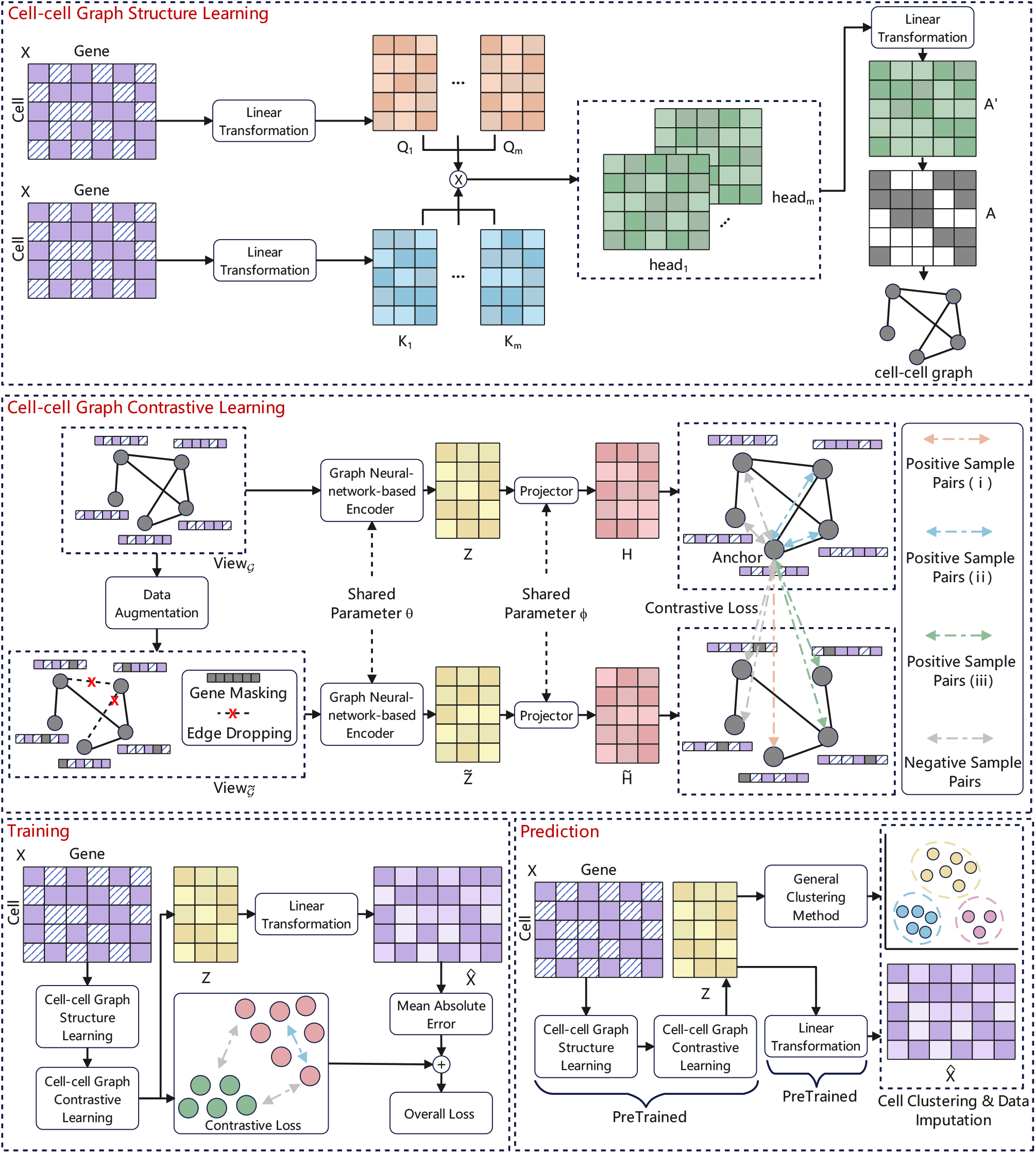
The overall framework of scSimGCL, which includes cell-cell graph structure and contrastive learning that take into account the contributions of cell-cell graph representations for cell clustering.

### Cell-cell graph structure learning

Let *𝒢* = (*𝒱, ℰ*, X) = (A, X) be a cell-cell graph. *V* = *{v*_1_, *v*_2_, *…, v*_*N*_ *}* is a collection of nodes, where each is a cell and *N* is the number of cells. ℰ is a collection of edges between nodes. X ∈ ℝ ^*N ×g*^ is the gene expression matrix obtained after preprocessing, where *g* is the number of genes. The genes with zero values in *X* are denoted by using a mask matrix M ∈ ℝ^*N×g*^. The aim of cell-cell graph structure learning is to obtain an adjacency matrix *A* ∈ {0, 1} ^*N ×N*^ using scRNA-seq data, where *A*_*ij*_ denotes the edge between cells *i* and *j*. In order to obtain the adjacency matrix *A*, we incorporate a multi-head attention module [40] into the network architecture. Specifically, query *Q* and key *K* vectors are generated using a linear transformation on *X*. The dot product is carried out between *Q* and *K*, using the Softmax function to obtain up to *m* attention heads. All heads are concatenated and then projected using a linear transformation to the original space.

The mathematical formulation can be written as follows:

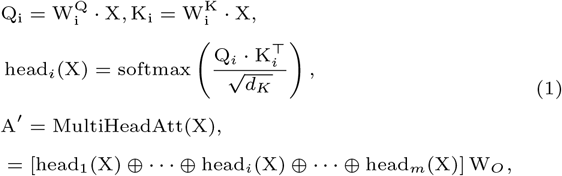

where *head*_*i*_ is the i-th attention head. *⊕* is the concatenation. *d*_*K*_ is the dimension of *K*. Accordingly, a sparse and non-negative adjacency matrix *A* can be obtained using the intermediate matrix A^*′*^ with numerical constraints as:

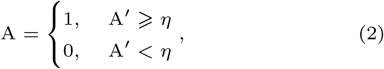

where *η* is a learnable threshold that excludes the values lower than that. Note that the proposed scSimGCL is an end-to-end cell representation learning framework where cell-cell graph structure learning and cell-cell graph contrastive learning work closely together. Accordingly, the proposed cell-cell graph structure learning module is not trained independently of the scSimGCL.

### Cell-cell graph contrastive learning

Data augmentation techniques allow the construction of counterparts from data samples. A data augmentation module is an important component in the GCL paradigm and plays a vital role in increasing the diversity of input graph samples [57]. We adopt attribute/gene masking and edge dropping [26] in the learned cell-cell graph to generate its counterpart. Specifically, we implement edge dropping by randomly dropping a collection of edges from the cell-cell graph. Given the adjacency matrix *A*, a masking matrix M^(*a*)^ ∈ {0, 1}^*N×N*^ is used here, where each 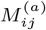 is derived from a Bernoulli distribution with probability *p*^(*a*)^ independently. Accordingly, a new Ã can be derived from using *A* with *M* as:

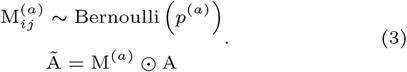

We implement gene masking by randomly masking a collection of genes with zero values from the cell-cell graph. In the same vein, a random vector M^(*x*)^ ∈ {0, 1}^*g*^ is used here, where each is derived from a Bernoulli distribution with probability *p*^(*x*)^ independently. Subsequently, the gene vector of each cell is masked with M^(x)^ and the gene matrix 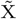 is generated as:

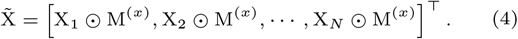

The outputs of the above process are Ã and 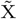. The two cell-cell graphs *𝒢* = (*𝒜*, X) and 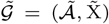 can be regarded as two graph views *V iew*_*𝒢*_ and 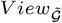 for contrastive learning. Before implementing contrastive learning, we employ a graph neural network-based encoder *f*_*θ*_(·) to generate node/cell representations as:

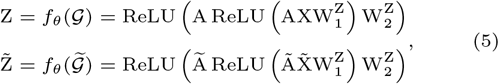

where *θ* is the parameter of *f* _*θ*_ (·). *Z* and 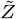 are cell representations for both *V iew*_*𝒢*_ and 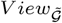 The graph neural network-based encoder employed here is graph convolutional networks [18]. Subsequently, a projector *f*_*ϕ*_(·) is used to map the generated cell representations Z and 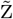 into a lower-dimensional latent space as:

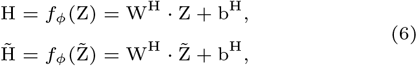

where *ϕ* is the parameter of *f*_*ϕ*_(·). *H* and 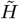 are cell representations obtained after projection. The projector used here is built upon a natural Multi-Layer Perceptrons. The cell representations *H* and 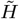 obtained through the above steps are used for contrastive learning.

Contrastive learning aims to maximize *H* and 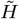 mutual information. A node/cell is arbitrarily chosen from the learned cell-cell graph view as an anchor. The positive sample pairs for contrastive learning are as follows: (i) the nodes connected to the anchor, (ii) the anchor counterpart in its counterpart view, and (iii) the nodes connected to the anchor from its counterpart view, while negative samples are the remaining samples. The i-th cell in *V iew*_*𝒢*_ is taken as the anchor and thus the contrastive loss 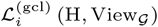 as:

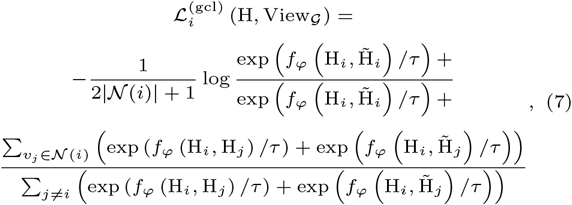

where *𝒩* (*i*) is a collection of neighbors for the i-th cell in *𝒢. f*_*φ*_(·) is the measurement of similarity, and the dot product is carried out here *f*_*φ*_(*a, b*) = *a* · *b. τ* is a temperature parameter that provides the penalty for negative pairs.

Since we have two graph views *V iew*_*𝒢*_ and 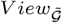 used for contrastive learning, the i-th cell in 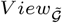 is chosen as an anchor again. The contrastive loss 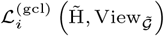 is computed by the way used in Eq. (7). Accordingly, the contrastive loss between *H* and 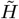 as:

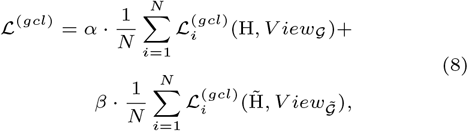

where *α* and *β* are two scaling parameters that take into account the contributions of the two losses.

### Training and Prediction

We design a pretraining mechanism to explore the topological nuances of cell-cell graphs. This mechanism implements contrastive learning with data imputation in a joint learning strategy. We have obtained contrastive learning loss *ℒ* ^(*gcl*)^ from the above section. Now, the imputation loss *ℒ* ^*(imp)*^ needs to be formulated. The cell representation *Z* for *V iew*_*𝒢*_ is used for scRNA-seq data imputation as:

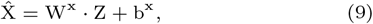

where 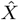 is the obtained scRNA-seq data after predicting. The imputation of scRNA-seq data is a regression task, and thus the mean absolute error between *X* and 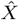 of each cell as:

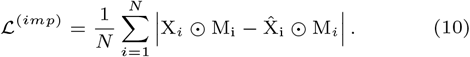

Next, we integrate both *ℒ* ^(*gcl*)^ and *ℒ* ^(*imp*)^ losses into an overall loss as:

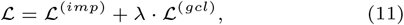

where *λ* is a scaling parameter that takes into account the contributions of *ℒ* ^(*imp*)^ and *ℒ* ^(*gcl*)^.

In summary, through the obtained cell representation *Z*, a general clustering method can be employed to carry out clustering assignments. Accordingly, the two stages, cell representation learning and clustering assignment analysis, are decoupled in the proposed scSimGCL framework. By doing this, our scSimGCL framework provides more flexibility in implementing different clustering algorithms (K-Means and Affinity Propagation).

### Data

We evaluate the proposed scSimGCL against competing deep learning approaches on simulated and real scRNA-seq datasets. We compare scSimGCL with state-of-the-art models on scRNA-seq data imputation and cell clustering tasks. The ten scRNA-seq datasets used are Shekhar mouse retina cells [32], Baron [2], 10X PBMC [58], Camp [3], Mouse bladder cells [13], Zeisel [55], Tabula Sapiens - Heart [8], Tabula Sapiens - Fat [8], Tabula Sapiens - Tongue [8], and Chien [5]. Each dataset is a two-dimensional matrix, where rows and columns are cells and genes, and each cell has its own available label. These datasets are processed using the Scanpy toolkit [45]. It includes excluding genes with expression values of zero from all cells first and then performing the normalization and logarithmically transformation on the gene expression matrix. Note that scSimGCL is not an approach proposed for batch effects. For example, when analyzing the Shekhar mouse retina cells dataset, we used all the data and did not discard any batches. The results obtained from the preliminary analysis of the ten datasets are set out in Table 1.

**Table 1.** Results of the statistical analysis on the ten datasets.

| Dataset | No. of cells | No. of genes | No. of cell types | Sequencing platform |
| --- | --- | --- | --- | --- |
| Shekhar mouse retina cells | 27499 | 13166 | 19 | Drop-seq |
| Baron | 8569 | 20125 | 14 | inDrop |
| 10X PBMC | 4340 | 33694 | 8 | 10x |
| Camp | 777 | 16270 | 7 | SMARTer |
| Mouse bladder cells | 2746 | 20670 | 16 | Microwell-seq |
| Zeisel | 3005 | 19972 | 9 | STRT-seq UMI |
| Tabula Sapiens - Heart | 10188 | 58482 | 5 | 10x 3' v3 |
| Tabula Sapiens - Fat | 19612 | 58482 | 12 | 10x 3' v3 |
| Tabula Sapiens - Tongue | 13629 | 58482 | 12 | 10x 3' v3 |
| Chien | 37121 | 60448 | 15 | mCT-seq |

The simulated scRNA-seq data is derived from the implementation of the Splatter [54] toolkit. The Splatter package is designed for simulating scRNA-seq data, offering a user-friendly interface for creating reproducible simulations. It allows the estimation of parameters from real data and provides functions for comparing real and simulated datasets. There are up to twelve simulated datasets created for three case studies. This includes the simulation of (i) the dropout events in the gene expression matrix, (ii) the strength of low signals for clustering assignments, and (iii) the high dimensionality of the gene expression matrix under extremely sparsity space. The three case studies are designed according to the previous study [35] and de.fracScale, dropout.mid, and dropout.shape mentioned below are all parameters of the Splatter package. In order to implement the case study with varied dropout rates, we set the value of dropout.mid from 0 to 2, i.e., 0.0, 0.5, 1.0, 1.5, and 2.0. Meanwhile, the number of cells is set to 6000 with up to 4 clusters, the number of genes is set to 5000, and dropout.shape and de.fracScale are set to -1 and 0.3, respectively. In order to implement the case study with the different strengths of low signals (Sigma), we set the value of de.fracScale from 0.1 to 0.25, i.e., 0.1, 0.15, 0.2, and 0.25. Meanwhile, the number of cells is set to 6000 with up to 4 clusters, the number of genes is set to 5000, and dropout.shape and dropout.mid are set to -1 and 0, respectively. In order to implement the case study with varied dimension sizes under extremely sparsity space, we set the number of genes from 10000 to 20000, i.e., 10000, 15000, and 20000, and the value of dropout.mid to 2. Meanwhile, the number of cells is set to 6000 with up to 4 clusters, the number of genes is set to 5000, and dropout.shape and de.fracScale are set to -1 and 0.3, respectively.

### Implementation and parameters setting

The proposed scSimGCL is developed using Python 3.8.17 with PyTorch 1.8.1. The scRNA-seq data are divided into two groups for analysis: 80% for model development and 20% for testing. For Cell-cell Graph Structure Learning, the number of attention heads *m* is 5, and the initial value of the learnable threshold *η* is 0.5. For Cell-cell Graph Contrastive Learning, the probability of a Bernoulli distribution is 0.5; the node size of the graph neural network-based encoder is 275; the node size of the projector is 80; the temperature parameter *τ* is 0.4, and the two scaling parameters *α* and *β* are 0.6 and 0.75, respectively. For Training and Prediction, the scaling parameter *λ* is 0.59. The Adam optimizer with a learning rate 1e-3 is employed to train the proposed approach. We utilize the default values of K-Means in Sklearn and specify the corresponding number of classes for each dataset. For other clustering algorithms, we utilize the default values provided in the relevant packages. The batch size for training depends on the sample size of the datasets. Specifically, a batch size of 128 is applied for datasets with a sample size below 4000, a batch size of 512 is applied for those with a sample size between 4000 and 8000, and a batch size of 1024 is applied for datasets exceeding 8000 samples. All parameters are obtained using the Grid Search method. The training is performed on a machine with a CPU: Intel Xeon Silver 4210, 256GB RAM, and GPU: Nvidia Titan RTX.

### Evaluation

The clustering accuracy (CA) [49], normalized mutual information (NMI) [33], and adjusted rand index (ARI) [17] are utilized to assess the performance of clustering assignments. Accordingly, the CA between the predicted cell type *Ŷ* with *N* _*Ŷ*_ clusters and the ground truth *Y* with *N*_*Y*_ clusters as:

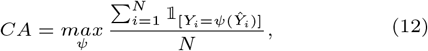

where *N* is the number of cells. *ψ* is a function that makes a transition between *Ŷ* and *Y* · 𝟙_[·]_ is an indicator function. The NMI between *Ŷ* and *Y* as:

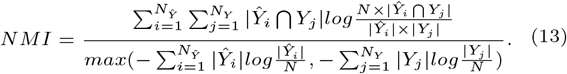

The ARI between *Ŷ* and *Y* as:

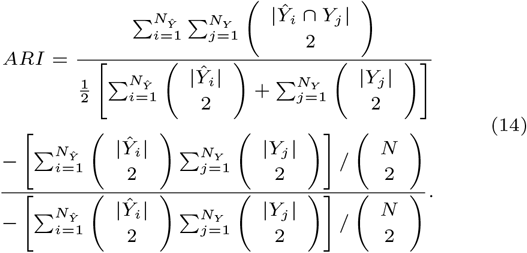

The Pearson correlation coefficients (PCCs) and L1 distance are employed to assess the performance of scRNA-seq data imputation as:

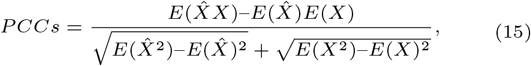

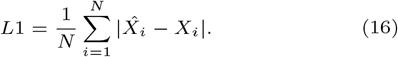

where *E*(·) is the expected value. 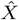 is the imputed value. *X* is the ground truth.

## Results and discussion

### Performance comparison with baselines

We evaluate our scSimGCL against competing clustering approaches on ten real scRNA-seq datasets. The clustering approaches employed in this study are scDCCA [42], scGCC [34], scGNN [43], graph-sc [7], CIDR [24], SIMLR [41], K-Means, Leiden [37], and Louvain [20]. The scDCCA is a combination of an autoencoder module and a contrastive learning module and focuses particularly on cell clustering. The scGCC comprises two modules, including a representation learning module and a clustering module and focuses particularly on cell clustering. The scGNN comprises three stacked autoencoders and focuses on the provision of scRNA-seq data imputation and cell clustering. The graph-sc is built upon a graph autoencoder and focuses particularly on cell clustering. The CIDR is a statistical model that includes reducing the impact of the high dimensionality of scRNA-seq data using principal component analysis on cell clustering. The SIMLR is built upon multi-kernel learning that performs dimension reduction and cell clustering by measuring cell similarity in scRNA-seq data. Louvain and Leiden are hierarchical clustering algorithms. Louvain merges communities into single nodes and performs modular clustering on compressed graphs, while Leiden is an improvement of Louvain and solves the problem of disconnected communities. We run all methods ten times and present the average CA, NMI, and ARI scores.

Figure 2 compares the CA, NMI, and ARI scores for each approach. We observed that the proposed scSimGCL consistently achieves the competing CA, NMI, and ARI scores compared with baselines. Significant improvement in our scSimGCL was found compared with K-Means. This result verifies the efficacy of our network architecture, which can generate high-quality representations and improve the performance of cell clustering. Looking at Figure 2, scDCCA is the best baseline for both Shekhar mouse retina cells and Zeisel datasets. Interestingly, the K-Means was observed to achieve the competing CA, NMI, and ARI scores on the Camp dataset compared to other baselines. This result may partly be explained by the fact that the performance of deep learning-based models is influenced by the input data size. As can be seen from Table 1, the Camp dataset contains a small sample size among the ten real scRNA-seq datasets. We run all methods ten times and present the average CA, NMI, and ARI scores.

**Fig. 2.**
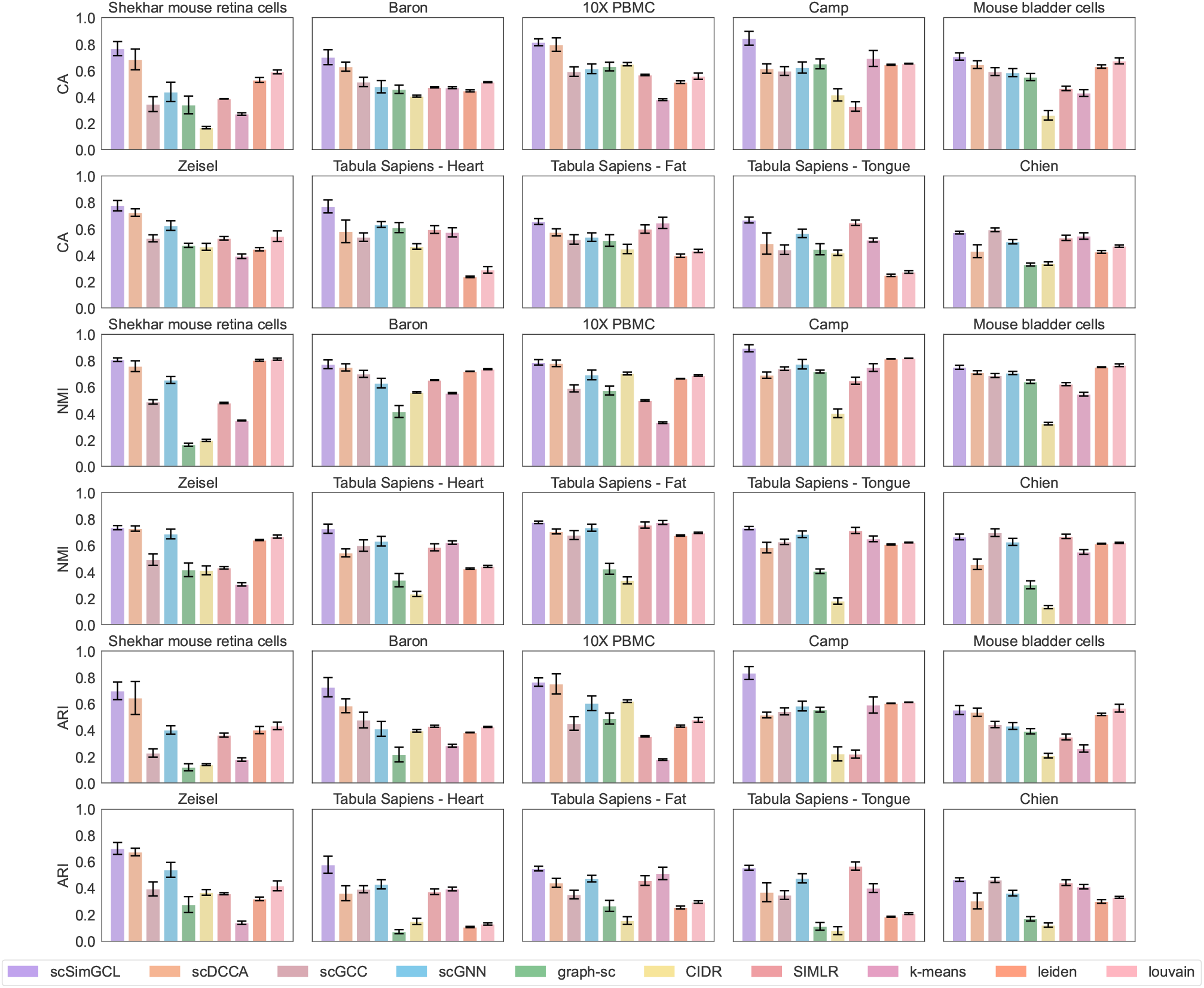
Clustering performance comparison using CA, NMI, and ARI on ten real scRNA-seq datasets. Higher values indicate better performance.

Figure 3 compares the results obtained from the clustering assignment of all baselines and our scSimGCL on twelve simulated scRNA-seq datasets. These results are average values obtained by running all approaches ten times. Our scSimGCL consistently achieves the competing CA, NMI, and ARI scores compared with baselines. After zooming on the subgraphs with varied dropout rates, we observed that there is an overlap between graph-sc and our scSimGCL. Thus, graph-sc is the best baseline among deep learning-based models, and it reported significantly more CA, NMI, and ARI scores than the other approaches. Moreover, graph-sc was observed to achieve the best accuracy in the case study with the different strengths of low signals. Despite its efficacy, there is a significant difference between graph-sc and our scSimGCL, which was evident in the data of de.fracScale 0.1 and 0.15. After zooming on the subgraphs with different dimension sizes, no difference in CA, NMI, and ARI scores of K-Means was observed. This rather intriguing result might be explained by the fact that K-Means lose its power as the number of dimensions increases. Together, these results provide important insights into the flexibility of deep learning-based models, i.e., clustering performance is superior to statistical models under extreme learning settings.

**Fig. 3.**
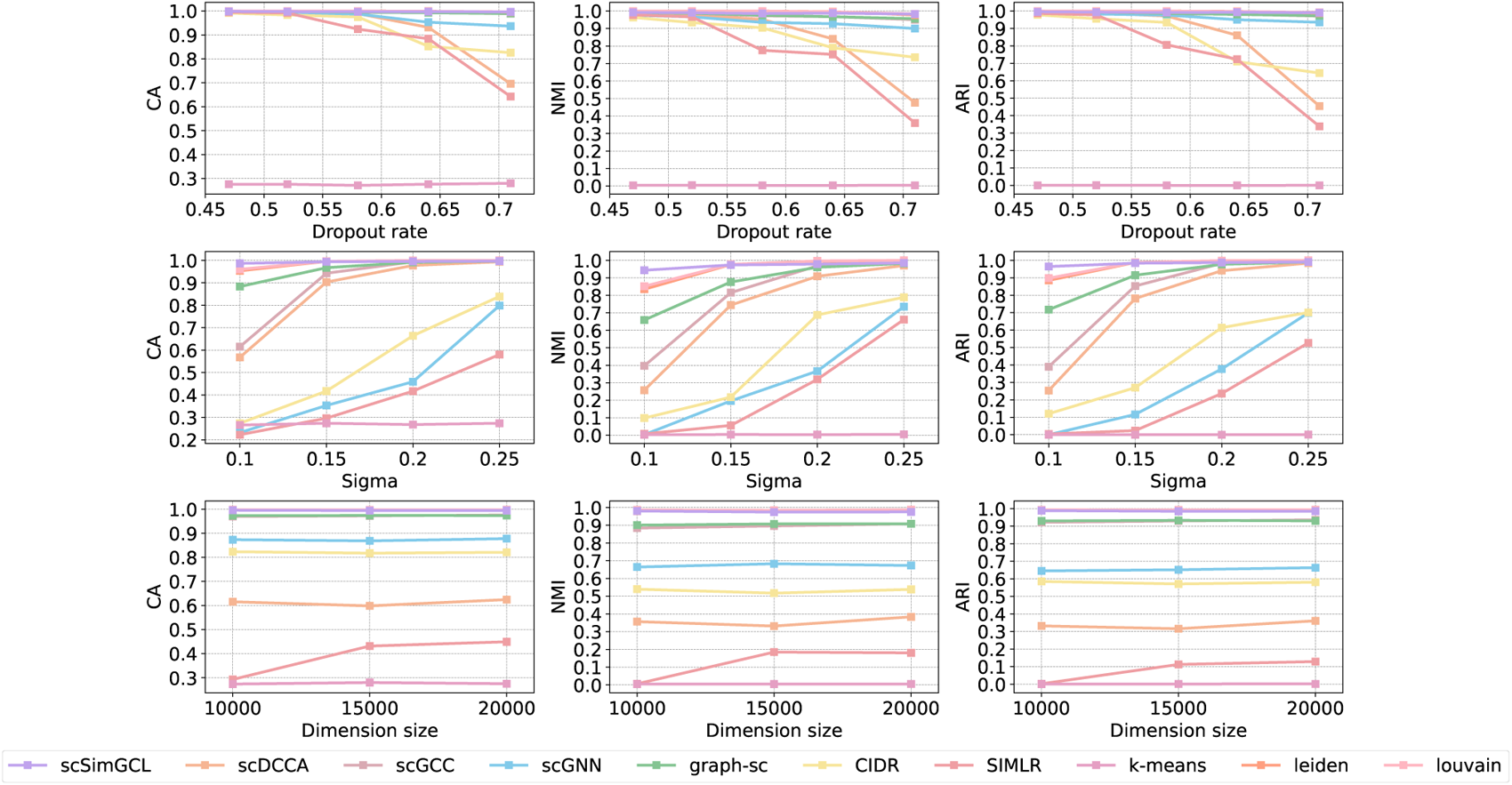
Clustering performance comparison using CA, NMI, and ARI on twelve simulated scRNA-seq datasets. Higher values indicate better performance.

Figure 4 displays the results obtained from the visualization analysis of all baselines and our scSimGCL on the Baron and Zeisel datasets. Specifically, the following steps should be done for Figure 4: the intermediate representation is obtained from each model and fed into the UMAP method [28] to generate a 2D embedding; the raw data is directly fed into the UMAP method to generate a 2D embedding. The difference between scSimGCL and baselines was significant. The identified clusters by our proposed scSimGCL were relatively visible compared to baselines. These findings enhance our understanding of model decisions, i.e., the transparency of deep learning models. However, caution must be applied with these identified clusters, as the findings might not be extrapolated to all datasets and learning settings. For example, the identified clusters by deep learning-based models are also known as intermediate representations and are therefore largely affected by model parameters or datasets. This has been illustrated by the results of scGNN on the two datasets. A note of caution is due here the input used by Leiden and Louvain in Figures 2, 3, and 4 is scRNA-seq data.

**Fig. 4.**
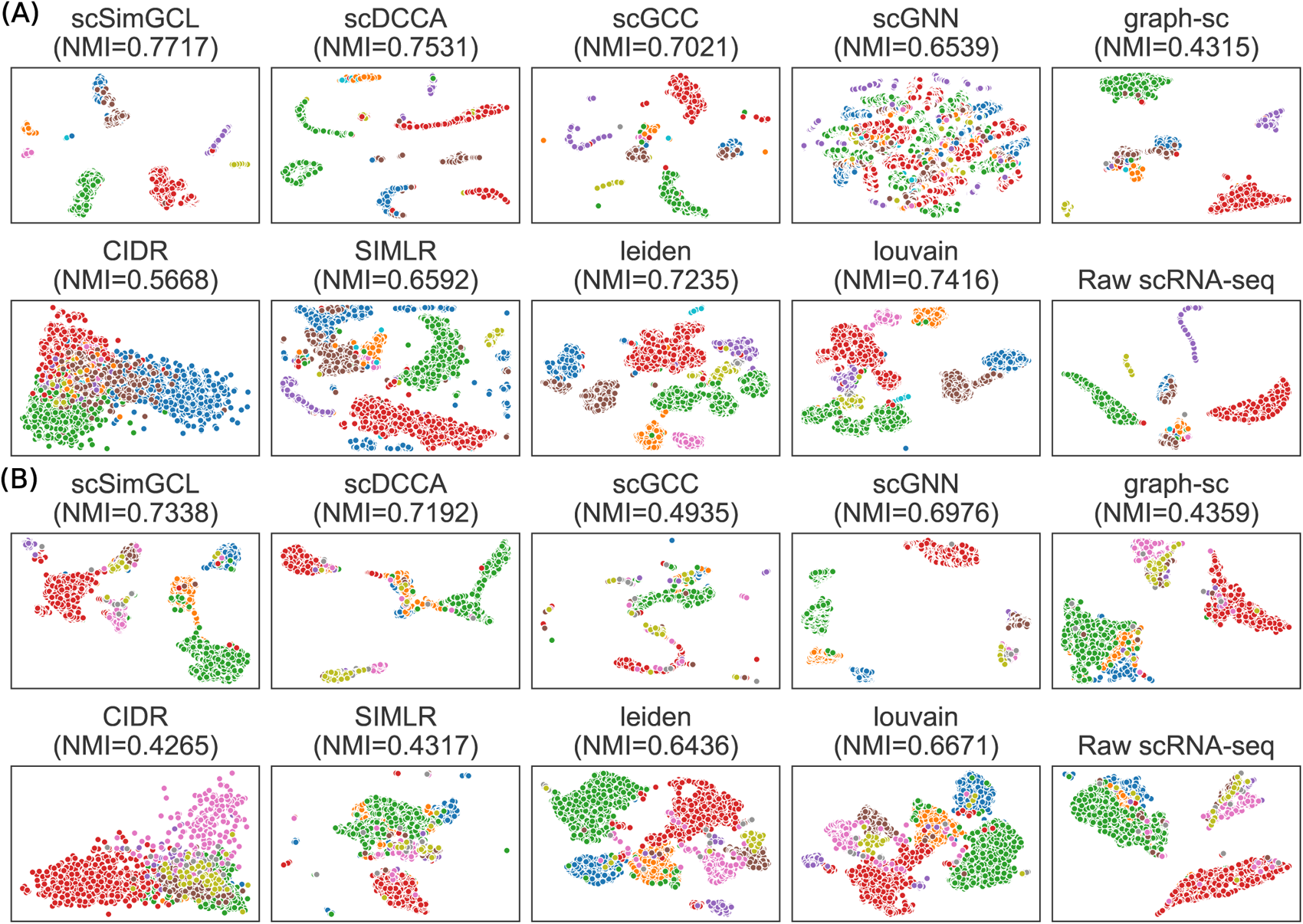
Visualization analysis results of baselines and scSimGCL on the (A) Baron and (B) Zeisel datasets.

Figure 5 displays the results of clustering algorithms on ten real scRNA-seq datasets. These results are average values obtained by running all approaches ten times. We feed the cell representation *Z* into the clustering algorithms used. Our scSimGCL is implemented with K-Means. K-Means, BIRCH, Leiden, Louvain, and Agglomerative Clustering can make decisions by indicating a predefined number of clusters. In contrast, Affinity Propagation can make decisions by automatically determining the number of clusters. The results of Louvain in the Shekhar mouse retina cells dataset are optimal. There was a significant difference between the performance of Affinity Propagation and other methods. This has been largely illustrated by the weak performance of Affinity Propagation on the four datasets, including Tabula Sapiens - Heart, Tabula Sapiens - Fat, Tabula Sapiens - Tongue, and Chien. The most interesting aspect of this graph is the competitive performance of Agglomerative Clustering on the Camp dataset. These results may be due to the size of the input data.

**Fig. 5.**
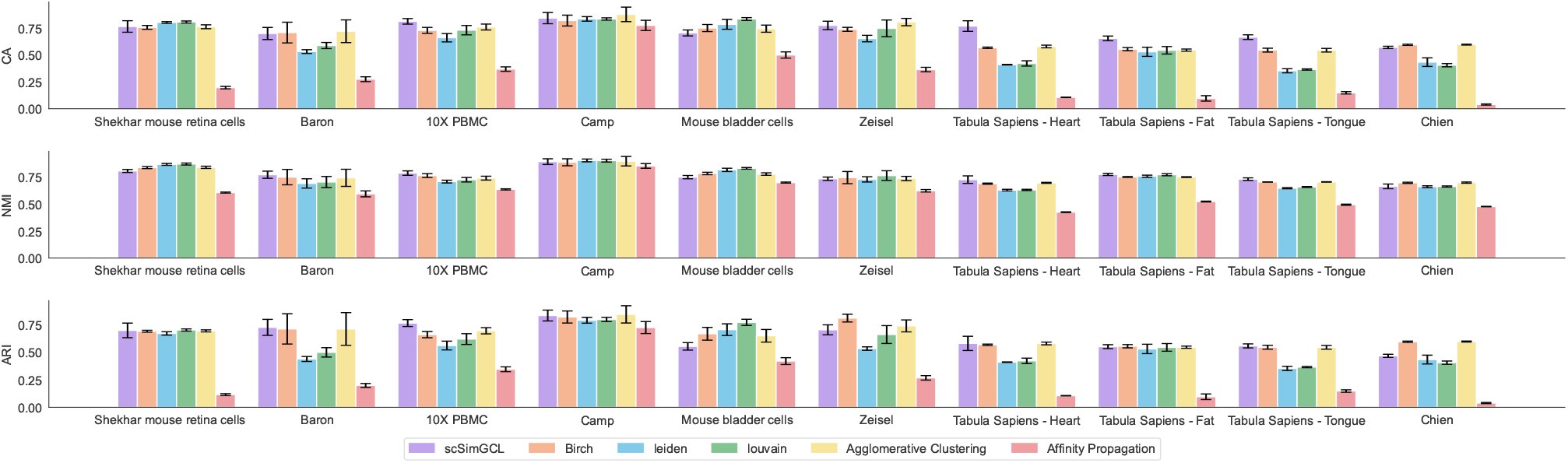
Performance comparison results of clustering algorithms on ten real scRNA-seq datasets using CA, NMI, and ARI. Higher values indicate better performance.

We evaluate our scSimGCL against competing scRNA-seq data imputation approaches on ten real scRNA-seq datasets. The scRNA-seq data imputation approaches employed in this study are GE-Impute [48], scGNN [43], scGCL [50], AutoClass [21], MAGIC [39], SAVER [16], and scImpute [22]. The GE-Impute is built upon graph neural networks and focuses particularly on scRNA-seq data imputation. Notably, GE-Impute incorporates the cell-cell similarity calculation into the imputation process, which is a crucial factor for improving imputation performance. The scGCL is a combination of graph contrastive learning and Zero-inflated Negative Binomial distribution and focuses particularly on scRNA-seq data imputation. The AutoClass comprises two neural networks, including an autoencoder and a classifier, which focuses on the provision of scRNA-seq data imputation and cell clustering. The MAGIC is a Markov affinity-based graph imputation of cells that includes a low-rank assumption for data propagation. Accordingly, MAGIC can retain data points in the low-frequency space and filter out noise/dropout points in the high-frequency space. The SAVER is a negative binomial model that imputes dropout values by estimating the distribution of input scRNA-seq data. The scImpute is a statistical model that focuses particularly on scRNA-seq data imputation. The core idea of scImpute is similar to GE-Impute, which imputes dropout values in the gene expression matrix by computing cell-cell similarity. We run all methods ten times and present the average PCCs score.

Figure 6 compares the PCCs and L1 distance for each approach. We observed that the proposed scSimGCL consistently achieves the competing PCCs and L1 distance compared with baselines. Looking at Figure 6, it is apparent that AutoClass is the best baseline for the Camp dataset. Besides, we observed that scGNN, scGCL, and AutoClass achieve lower PCCs but higher L1 distance on the Baron and Zeisel datasets. The performance variations of deep learning-based models are interesting but not surprising. This result may have something to do with the dropout rate of the input data or the sensitivity and specificity of deep learning-based models.

**Fig. 6.**
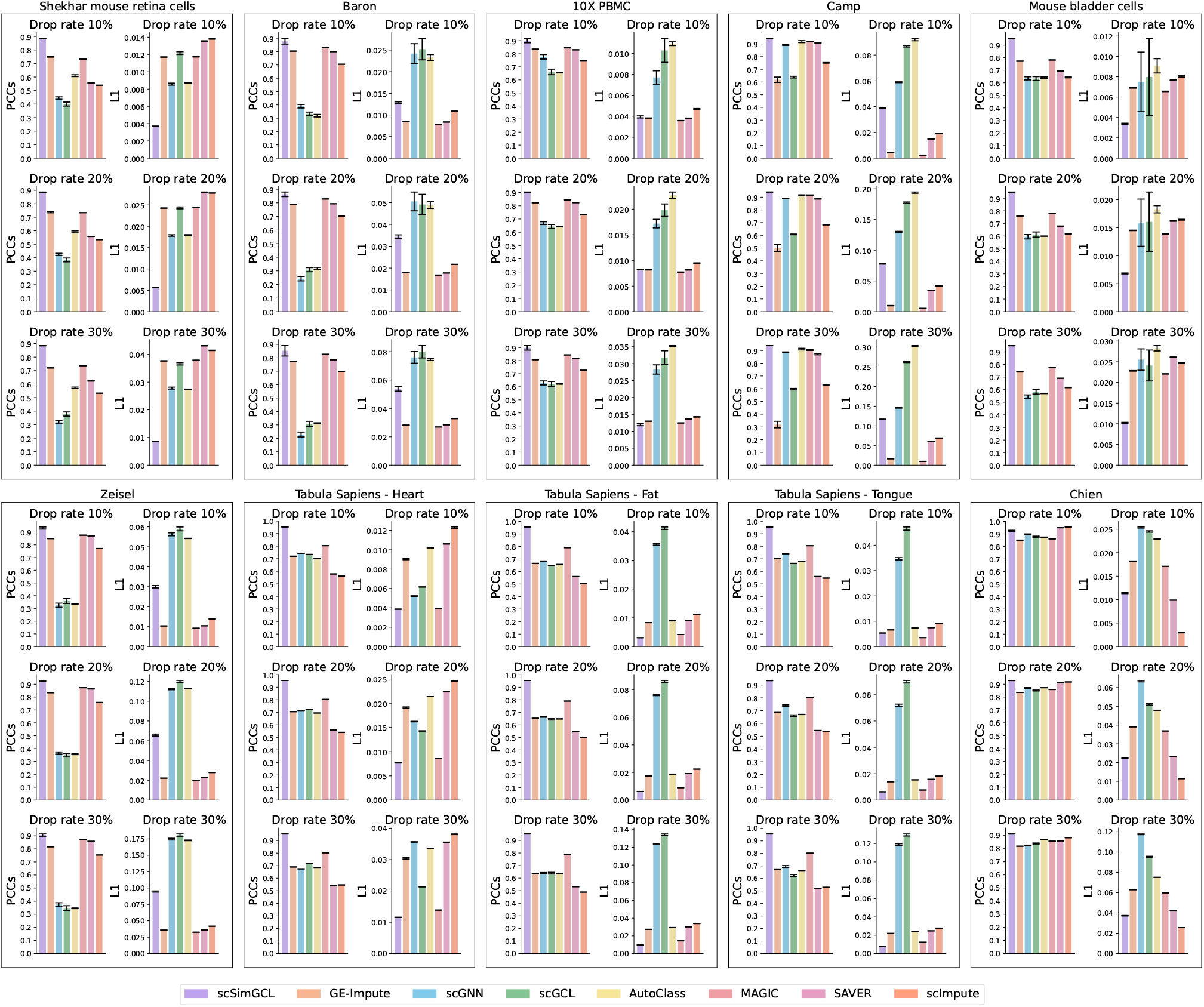
Imputation performance comparison on ten real scRNA-seq datasets using PCC and L1 distance. The higher the PCC value, the better the performance. The lower the L1 distance value, the better the performance.

### Ablation study

We propose three variants of scSimGCL to determine the efficacy of the designed GCL framework. These three variants are termd as scSimGCL_*α*_, scSimGCL_*β*_, and scSimGCL_*γ*_. In particular, the positive samples for the anchor used in scSimGCL_*α*_ are the nodes connected to the anchor and the nodes connected to the anchor from its counterpart view; the positive samples for the anchor used in scSimGCL_*β*_ are the nodes connected to the anchor and the anchor counterpart in its counterpart view; the positive samples for the anchor used in scSimGCL_*γ*_ are the anchor counterpart in its counterpart view. We run all methods ten times and present the average CA, NMI, ARI, PCCs, and L1 distance. The results of the ablation study are summarized in Figure 7A. On average, scSimGCL was shown to have achieved the most satisfactory performance on cell clustering and scRNA-seq data imputation. These results provide further support for the homophily assumption in the graph, which can facilitate the formation of informative positive and negative sample pairs, thereby improving the performance of graph contrastive learning.

**Fig. 7.**
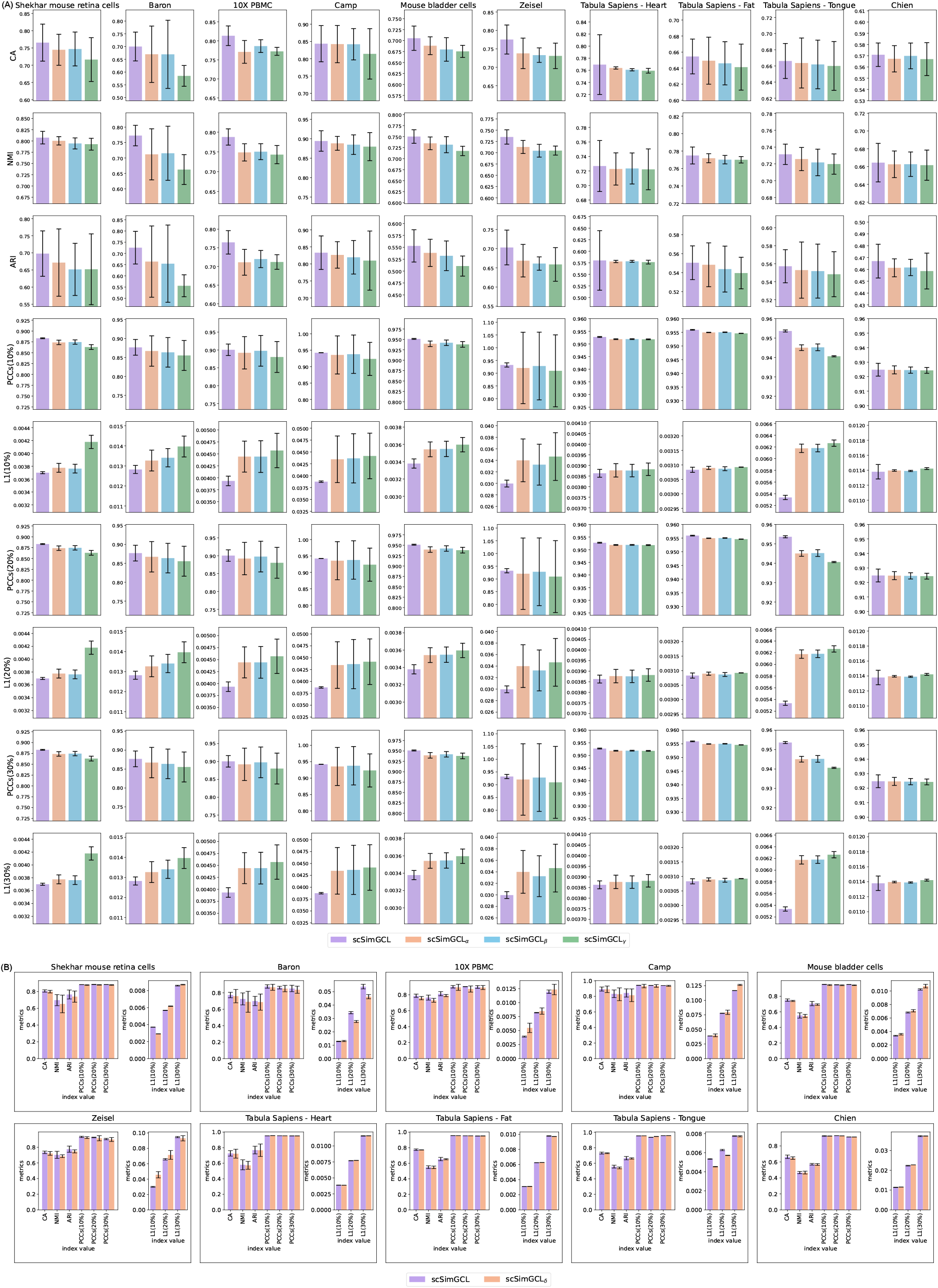
Performance comparison results between scSimGCL and its four variants.

We propose a variant of scSimGCL to determine the robustness of decisions in the self-supervised learning setting. The proposed scSimGCL_*δ*_ includes two implementation settings, where *ℒ*^(*gcl*)^ and *ℒ*^(*imp*)^ are separately utilized as the sole objective function in cluster assignment and data imputation. Note that the percentages used in PCCs (10%), PCCs (20%), PCCs (30%), L1 (10%), L1 (20%), and L1 (30%) are dropout rates. We run all methods ten times and present the average CA, NMI, ARI, PCCs, and L1 distance. Figure 7B compares CA, NMI, ARI, PCCs, and L1 distance for scSimGCL and scSimGCL_*δ*_. Compared with the data in scSimGCL, the performance of scSimGCL_*δ*_ on both cluster assignment and data imputation has a clear decreasing trend. Accordingly, contrastive learning and data imputation in the designed pretraining mechanism are inseparable, which is helpful for improving the overall performance of our network architecture.

### Analysis of Hyperparameters

We present a hyperparameter analysis to determine the efficacy of our network architecture. There are batch size, temperature *τ*, learnable threshold *η*, dropout rate *p*^(*drop*)^, and scaling parameter *λ* that we are interested in. A note of caution is due here since the dropout rate in neural networks differs from that in scRNA-seq datasets. We implement the dropout rates in neural networks with the obtained cell representation *Z*. Batch size and dropout rate have long been a question of great interest in neural networks. Temperature parameter is often used in research on contrastive learning. Scaling parameter is used to make tradeoff the contributions of ℒ^(*imp*)^ and *ℒ*^(*gcl*)^. Note that varied batch sizes are implemented using three datasets with a relatively large sample size, as shown in Figure 8. These results are average values obtained by running scSimGCL with different parameters ten times. After zooming on the subgraphs with varied batch sizes (Figure 8A), we observed that PCCs in the three datasets could reach the optimal value on the same batch size. In the same vein, CA, NMI, and ARI in the Shekhar mouse retina cells and 10X PBMC datasets can reach optimal values on the same batch size. As can be seen from the subgraphs below (Figure 8B), our approach could make decisions based on varied parameters *ϕ, τ, p*^(*drop*)^, and *λ*. We have also examined the impact of *α* and *β* (i.e., two scaling parameters in Eq. (8)), *p*^(*a*)^ and *p*^(*x*)^ (i.e., edge dropping and gene masking) on the model decisions. The results, as shown in Figure 9, were obtained from the Zeisel dataset. This analysis was successful as it was able to identify patterns and trends in the data.

**Fig. 8.**
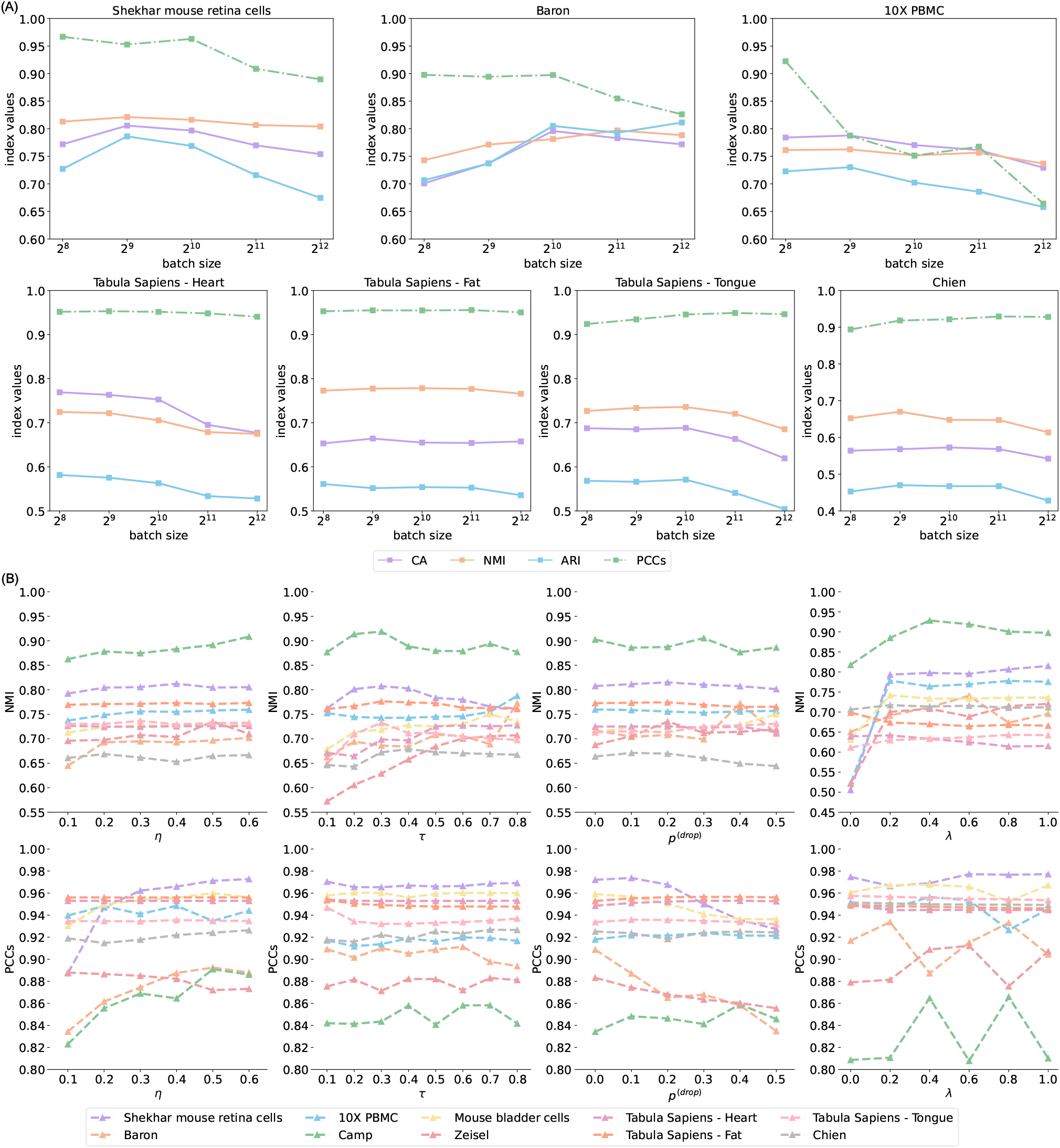
Hyperparameter analysis results using CA, NMI, ARI, and PCCs. The batch size, learnable threshold *η*, dropout rate *p*^(*drop*)^, and scaling parameter *λ* were carried out in the analysis. Higher values indicate better performance.

**Fig. 9.**
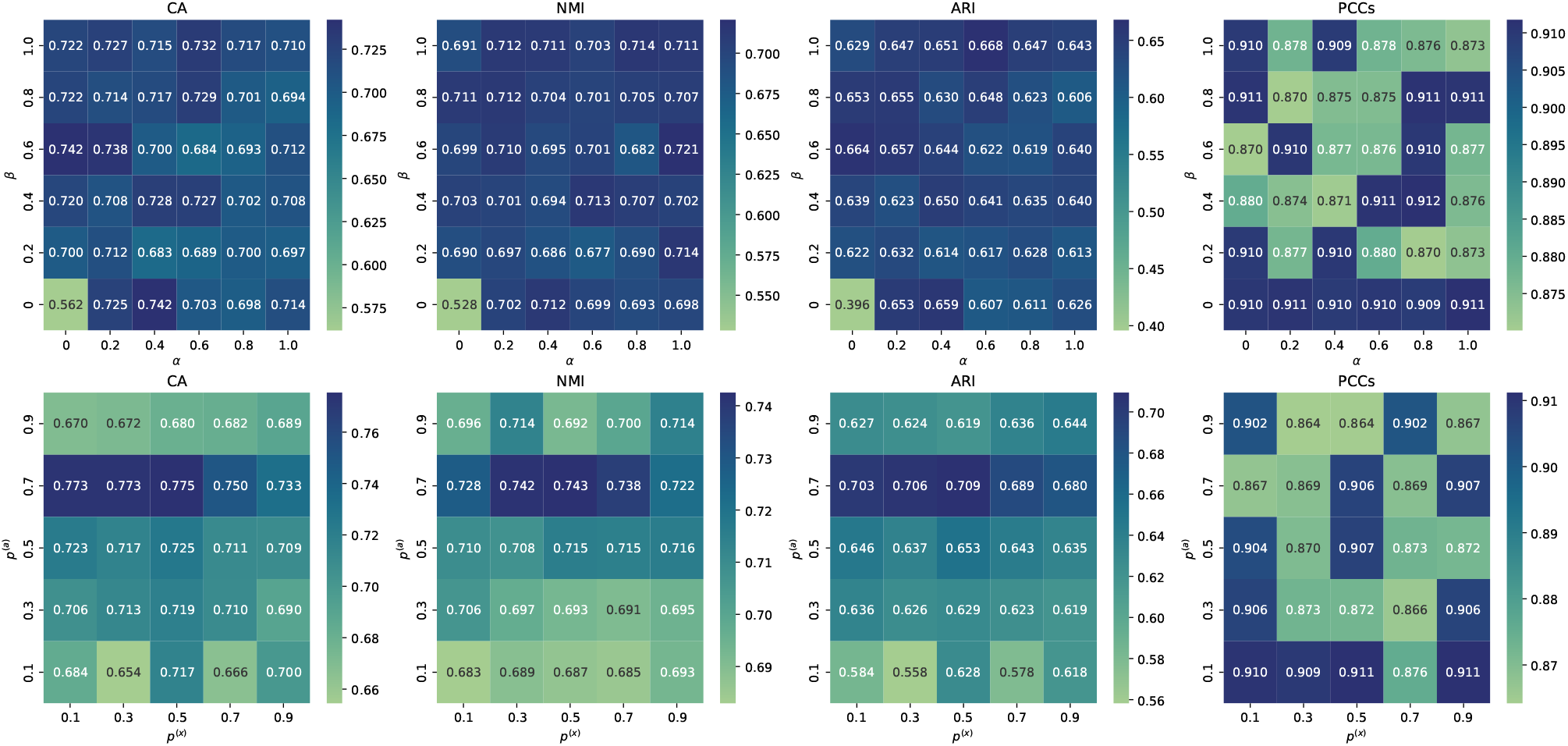
Hyperparameter analysis results using CA, NMI, ARI, and PCCs. The edge dropping *p*^(*a*)^ and gene masking *p*^(*x*)^ were carried out in the analysis. Higher values indicate better performance.

### Cell Trajectory Inference

Trajectory inference in scRNA-seq data analysis allows researchers to identify critical stages and transitions in cell development. We assess the effectiveness of the representations generated by scSimGCL using the Yan [51] dataset and focus on the developmental process of mouse embryos. The UMAP graph in Figure 10 is generated using the UMAP method [28]. It maps high-dimensional gene expression data into a 2D plane for visualization analysis. In particular, the representation generated from scSimGCL (*Z* in Equation 5) and the PAGA method [46] are used to generate cell developmental trajectories, as shown in the PAGA graph. We can see that the entire cell development trajectory starts from the zygote cell, passes through 2, 4, 8, and 16 cells, and ends with the blast cell. Overall, this result agrees with the findings of the actual mouse embryo development process. The findings in this trajectory inference are subject to one limitation: there is a disconnect between the 2-cell and 4-cell clusters.

**Fig. 10.**
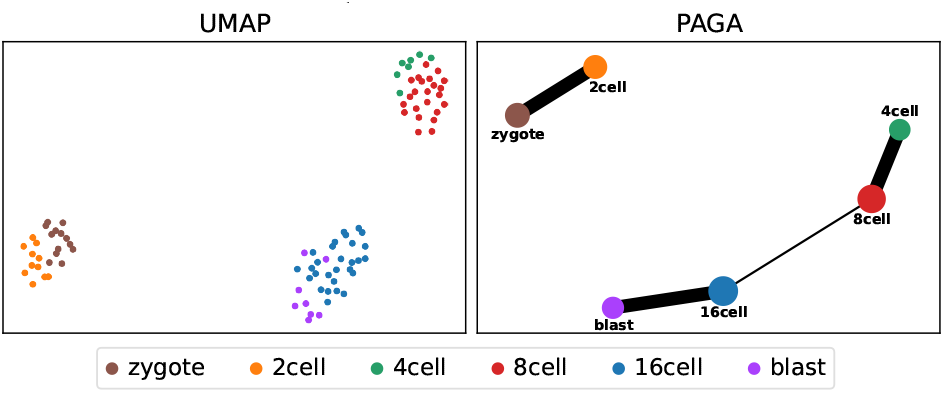
Cell trajectory inference on the Yan dataset.

### The run-time analysis of each model

Figure 11 compares the clustering and imputation performance on the Camp dataset and presents the training time of scSimGCL on ten datasets. From the graph below, we can see that the proposed scSimGCL achieved higher NMI and PCC scores. On average, the run time of K-Means, Leiden, and Louvain was lower than the deep neural network models such as scDCCA and scGNN. Interestingly, the MAGIC was observed to achieve the competing PCC scores, but the run time of MAGIC was lower than that of other models. It seems possible that these results are due to the computational complexity of deep learning, i.e., the computational complexity of deep neural network models is usually higher than that of traditional machine learning or statistical models.

**Fig. 11.**
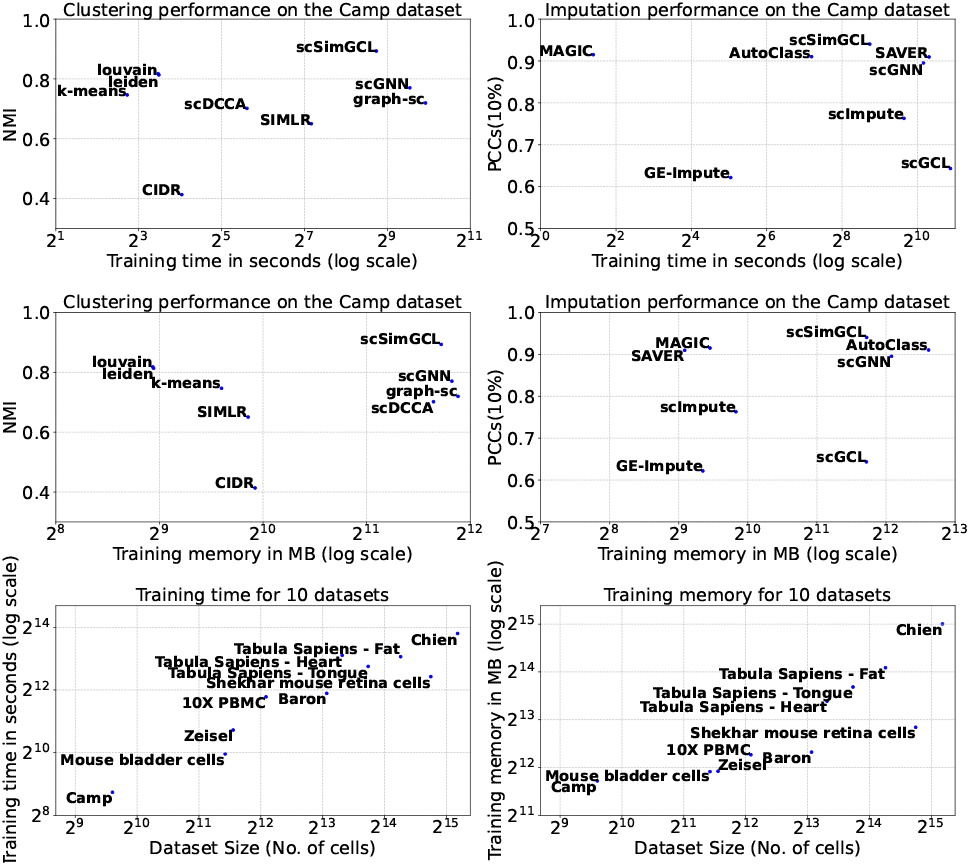
Clustering and imputation performance of baselines and our scSimGCL on the Camp dataset and the training time of scSimGCL on ten datasets.

## Conclusion

Single-cell RNA sequencing has become the most popular method for investigating transcriptome-wide gene expression at the single-cell level. Using scRNA-seq data on cell clustering is a well-established approach to provide a detailed understanding of the groups of cells with similar expression profiles. To achieve this aim, we need to solve challenges stemming from raw scRNA-seq data, including high dimensionality and dropout phenomenon. The existing deep learning-based solutions for addressing these challenges are extensive and focus particularly on graph machine learning models and contrastive learning-based models. There is still much work to be done to achieve desirable representations and accuracy simultaneously.

This study set out to marry graph neural networks with contrastive learning for scRNA-seq data analysis. We propose scSimGCL, a simple and effective framework based on the graph contrastive learning paradigm that leverages self-supervised pretraining of graph neural networks to generate high-quality representations critical for cell clustering.

Extensive experiments were carried out on simulated and real scRNA-seq datasets. Our scSimGCL demonstrated significant improvement in scRNA-seq data imputation and cell clustering. The results of clustering assignment analysis indicated that scSimGCL is a general approach that can include competing clustering algorithms. Ablation study and hyperparameter analysis further confirmed the efficacy of our network architecture with the robustness of decisions in the self-supervised learning setting.

With regard to the research method, a major limitation needs to be acknowledged here, i.e., the lack of uncertainty estimation in model decisions (associated with the over-confidence problem in deep learning approaches) adds caution regarding the generalizability of findings. Further modeling work will have to be conducted in order to improve the transparency and safety of scSimGCL, such as developing interpretable and reliable analytical approaches. Batch effects in scRNA-seq data may lead to false conclusions, as they arise when variations in the sample group are due to technical factors rather than biological realities. A further study with more focus on batch effects in scRNA-seq data is therefore suggested. Another possible area of future research would be to explore the potential use of scSimGCL on the scATAC-seq datasets. Since the dropout effects in scATAC-seq data are higher than those in scRNA-seq data, the new understanding should help assess the effectiveness and robustness of our scSimGCL.

## Data availability

The scRNAseq data that support the findings of this paper are publicly available at GitHub: https://github.com/zhangzh1328/scSimGCL.

## Code availability

The source code of scSimGCL is publicly available at https://github.com/zhangzh1328/scSimGCL.

### Key Points

- scSimGCL is a simple and effective graph contrastive learning framework for learning high-quality representations crucial for robust cell clustering.
- scSimGCL outperforms competing state-of-the-art models in scRNA-seq data imputation and cell clustering by a significant margin.
- scSimGCL is capable of making informed decisions by indicating a predefined number of clusters or automatically determining the number of clusters, which is a critical aspect in unsupervised learning settings for scRNA-seq data analysis.
- scSimGCL represents a further step towards developing a foundation model for cell clustering.

## References

1. Aisha A AlJanahi, Mark Danielsen, and Cynthia E Dunbar. An introduction to the analysis of single-cell rna-sequencing data. Molecular Therapy Methods & Clinical Development, 10:189–196, 2018.

2. Maayan Baron, Adrian Veres, Samuel L Wolock, Aubrey L Faust, Renaud Gaujoux, Amedeo Vetere, Jennifer Hyoje Ryu, Bridget K Wagner, Shai S Shen-Orr, Allon M Klein, et al. A single-cell transcriptomic map of the human and mouse pancreas reveals inter-and intra-cell population structure. Cell systems, 3(4):346–360, 2016.

3. J Gray Camp, Keisuke Sekine, Tobias Gerber, Henry Loeffler-Wirth, Hans Binder, Malgorzata Gac, Sabina Kanton, Jorge Kageyama, Georg Damm, Daniel Seehofer, et al. Multilineage communication regulates human liver bud development from pluripotency. Nature, 546(7659):533–538, 2017.

4. Yi Cheng and Xiuli Ma. scgac: a graph attentional architecture for clustering single-cell rna-seq data. Bioinformatics, 38(8):2187–2193, 2022.

5. Jo-Fan Chien, Hanqing Liu, Bang-An Wang, Chongyuan Luo, Anna Bartlett, Rosa Castanon, Nicholas D Johnson, Joseph R Nery, Julia Osteen, Junhao Li, et al. Cell-type-specific effects of age and sex on human cortical neurons. Neuron, 2024.

6. Madalina Ciortan and Matthieu Defrance. Contrastive self-supervised clustering of scrna-seq data. BMC bioinformatics, 22(1):280, 2021.

7. Madalina Ciortan and Matthieu Defrance. Gnn-based embedding for clustering scrna-seq data. Bioinformatics, 38(4):1037–1044, 2022.

8. The Tabula Sapiens Consortium*, Robert C Jones, Jim Karkanias, Mark A Krasnow, Angela Oliveira Pisco, Stephen R Quake, Julia Salzman, Nir Yosef, Bryan Bulthaup, Phillip Brown, et al. The tabula sapiens: A multiple-organ, single-cell transcriptomic atlas of humans. Science, 376(6594):eabl4896, 2022.

9. Gökcen Eraslan, Lukas M Simon, Maria Mircea, Nikola S Mueller, and Fabian J Theis. Single-cell rna-seq denoising using a deep count autoencoder. Nature communications, 10(1):390, 2019.

10. Yanglan Gan, Yuhan Chen, Guangwei Xu, Wenjing Guo, and Guobing Zou. Deep enhanced constraint clustering based on contrastive learning for scrna-seq data. Briefings in Bioinformatics, 24(4):bbad222, 2023.

11. Renxiang Guan, Zihao Li, Xianju Li, and Chang Tang. Pixel-superpixel contrastive learning and pseudo-label correction for hyperspectral image clustering. In ICASSP 2024-2024 IEEE International Conference on Acoustics, Speech and Signal Processing (ICASSP), pages 6795–6799. IEEE, 2024.

12. Renxiang Guan, Zihao Li, Wenxuan Tu, Jun Wang, Yue Liu, Xianju Li, Chang Tang, and Ruyi Feng. Contrastive multi-view subspace clustering of hyperspectral images based on graph convolutional networks. IEEE Transactions on Geoscience and Remote Sensing, 2024.

13. Xiaoping Han, Renying Wang, Yincong Zhou, Lijiang Fei, Huiyu Sun, Shujing Lai, Assieh Saadatpour, Ziming Zhou, Haide Chen, Fang Ye, et al. Mapping the mouse cell atlas by microwell-seq. Cell, 172(5):1091–1107, 2018.

14. Dayu Hu, Ke Liang, Zhibin Dong, Jun Wang, Yawei Zhao, and Kunlun He. Effective multi-modal clustering method via skip aggregation network for parallel scrna-seq and scatac-seq data. Briefings in Bioinformatics, 25(2):bbae102, 2024.

15. Dayu Hu, Ke Liang, Sihang Zhou, Wenxuan Tu, Meng Liu, and Xinwang Liu. scdfc: a deep fusion clustering method for single-cell rna-seq data. Briefings in Bioinformatics, 24(4):bbad216, 2023.

16. Mo Huang, Jingshu Wang, Eduardo Torre, Hannah Dueck, Sydney Shaffer, Roberto Bonasio, John I Murray, Arjun Raj, Mingyao Li, and Nancy R Zhang. Saver: gene expression recovery for single-cell rna sequencing. Nature methods, 15(7):539–542, 2018.

17. Lawrence Hubert and Phipps Arabie. Comparing partitions. Journal of classification, 2:193–218, 1985.

18. Thomas N Kipf and Max Welling. Semi-supervised classification with graph convolutional networks. arXiv preprint arXiv:1609.02907, 2016.

19. Junseok Lee, Sungwon Kim, Dongmin Hyun, Namkyeong Lee, Yejin Kim, and Chanyoung Park. Deep single-cell rna-seq data clustering with graph prototypical contrastive learning. Bioinformatics, 39(6):btad342, 2023.

20. Jacob H Levine, Erin F Simonds, Sean C Bendall, Kara L Davis, D Amir El-ad, Michelle D Tadmor, Oren Litvin, Harris G Fienberg, Astraea Jager, Eli R Zunder, et al. Data-driven phenotypic dissection of aml reveals progenitor-like cells that correlate with prognosis. Cell, 162(1):184–197, 2015.

21. Hui Li, Cory R Brouwer, and Weijun Luo. A universal deep neural network for in-depth cleaning of single-cell rna-seq data. Nature Communications, 13(1):1901, 2022.

22. Wei Vivian Li and Jingyi Jessica Li. An accurate and robust imputation method scimpute for single-cell rna-seq data. Nature communications, 9(1):997, 2018.

23. Xiangjie Li, Kui Wang, Yafei Lyu, Huize Pan, Jingxiao Zhang, Dwight Stambolian, Katalin Susztak, Muredach P Reilly, Gang Hu, and Mingyao Li. Deep learning enables accurate clustering with batch effect removal in single-cell rna-seq analysis. Nature communications, 11(1):2338, 2020.

24. Peijie Lin, Michael Troup, and Joshua WK Ho. Cidr: Ultrafast and accurate clustering through imputation for single-cell rna-seq data. Genome biology, 18:1–11 2017.

25. Tianxiang Liu, Yue Bi, Xudong Guo, Quan Zou, Cangzhi Jia, and Fuyi Li. scdfn: Enhancing single-cell rna-seq clustering with deep fusion networks. 2024.

26. Yixin Liu, Yu Zheng, Daokun Zhang, Hongxu Chen, Hao Peng, and Shirui Pan. Towards unsupervised deep graph structure learning. In Proceedings of the ACM Web Conference 2022, pages 1392–1403, 2022.

27. Yuxi Liu, Zhenhao Zhang, Shaowen Qin, Flora D Salim, and Antonio Jimeno Yepes. Contrastive learning-based imputation-prediction networks for in-hospital mortality risk modeling using ehrs. In Joint European Conference on Machine Learning and Knowledge Discovery in Databases, pages 428–443. Springer, 2023.

28. Leland McInnes, John Healy, and James Melville. Umap: Uniform manifold approximation and projection for dimension reduction. arXiv preprint arXiv:1802.03426, 2018.

29. Michael Moor, Oishi Banerjee, Zahra Shakeri Hossein Abad, Harlan M Krumholz, Jure Leskovec, Eric J Topol, and Pranav Rajpurkar. Foundation models for generalist medical artificial intelligence. Nature, 616(7956):259–265, 2023.

30. Cheng Peng, Xi Yang, Mengxian Lyu, Kaleb E Smith, Mona G Flores, Jiang Bian, and Yonghui Wu. Gatortron and gatortrongpt: Large language models for clinical narratives. In AAAI 2024 Spring Symposium on Clinical Foundation Models, 2024.

31. Peng Qiu. Embracing the dropouts in single-cell rna-seq analysis. Nature communications, 11(1):1169, 2020.

32. Karthik Shekhar, Sylvain W Lapan, Irene E Whitney, Nicholas M Tran, Evan Z Macosko, Monika Kowalczyk, Xian Adiconis, Joshua Z Levin, James Nemesh, Melissa Goldman, et al. Comprehensive classification of retinal bipolar neurons by single-cell transcriptomics. Cell, 166(5):1308–1323, 2016.

33. Alexander Strehl and Joydeep Ghosh. Cluster ensembles— a knowledge reuse framework for combining multiple partitions. Journal of machine learning research, 3(Dec):583–617, 2002.

34. Sheng-Wen Tian, Jian-Cheng Ni, Yu-Tian Wang, Chun-Hou Zheng, and Cun-Mei Ji. scgcc: Graph contrastive clustering with neighborhood augmentations for scrna-seq data analysis. IEEE Journal of Biomedical and Health Informatics, 2023.

35. Tian Tian, Ji Wan, Qi Song, and Zhi Wei. Clustering single-cell rna-seq data with a model-based deep learning approach. Nature Machine Intelligence, 1(4):191–198, 2019.

36. Tian Tian, Jie Zhang, Xiang Lin, Zhi Wei, and Hakon Hakonarson. Model-based deep embedding for constrained clustering analysis of single cell rna-seq data. Nature communications, 12(1):1873, 2021.

37. Vincent A Traag, Ludo Waltman, and Nees Jan Van Eck. From louvain to leiden: guaranteeing well-connected communities. Scientific reports, 9(1):1–12, 2019.

38. Bram Van de Sande, Joon Sang Lee, Euphemia Mutasa-Gottgens, Bart Naughton, Wendi Bacon, Jonathan Manning, Yong Wang, Jack Pollard, Melissa Mendez, Jon Hill, et al. Applications of single-cell rna sequencing in drug discovery and development. Nature Reviews Drug Discovery, 22(6):496–520, 2023.

39. David Van Dijk, Roshan Sharma, Juozas Nainys, Kristina Yim, Pooja Kathail, Ambrose J Carr, Cassandra Burdziak, Kevin R Moon, Christine L Chaffer, Diwakar Pattabiraman, et al. Recovering gene interactions from single-cell data using data diffusion. Cell, 174(3):716–729, 2018.

40. Ashish Vaswani, Noam Shazeer, Niki Parmar, Jakob Uszkoreit, Llion Jones, Aidan N Gomez, Łukasz Kaiser, and Illia Polosukhin. Attention is all you need. Advances in neural information processing systems, 30, 2017.

41. Bo Wang, Junjie Zhu, Emma Pierson, Daniele Ramazzotti, and Serafim Batzoglou. Visualization and analysis of single-cell rna-seq data by kernel-based similarity learning. Nature methods, 14(4):414–416, 2017.

42. Jing Wang, Junfeng Xia, Haiyun Wang, Yansen Su, and Chun-Hou Zheng. scdcca: deep contrastive clustering for single-cell rna-seq data based on auto-encoder network. Briefings in Bioinformatics, 24(1):bbac625, 2023.

43. Juexin Wang, Anjun Ma, Yuzhou Chang, Jianting Gong, Yuexu Jiang, Ren Qi, Cankun Wang, Hongjun Fu, Qin Ma, and Dong Xu. scgnn is a novel graph neural network framework for single-cell rna-seq analyses. Nature communications, 12(1):1882, 2021.

44. Shudong Wang, Yu Zhang, Yulin Zhang, Wenhao Wu, Lan Ye, YunYin Li, Jionglong Su, and Shanchen Pang. scasgc: An adaptive simplified graph convolution model for clustering single-cell rna-seq data. Computers in Biology and Medicine, 163:107152, 2023.

45. F Alexander Wolf, Philipp Angerer, and Fabian J Theis. Scanpy: large-scale single-cell gene expression data analysis. Genome biology, 19:1–5, 2018.

46. F Alexander Wolf, Fiona K Hamey, Mireya Plass, Jordi Solana, Joakim S Dahlin, Berthold Göttgens, Nikolaus Rajewsky, Lukas Simon, and Fabian J Theis. Paga: graph abstraction reconciles clustering with trajectory inference through a topology preserving map of single cells. Genome biology, 20:1–9, 2019.

47. Michael Wornow, Yizhe Xu, Rahul Thapa, Birju Patel, Ethan Steinberg, Scott Fleming, Michael A Pfeffer, Jason Fries, and Nigam H Shah. The shaky foundations of large language models and foundation models for electronic health records. npj Digital Medicine, 6(1):135, 2023.

48. Xiaobin Wu and Yuan Zhou. Ge-impute: graph embedding-based imputation for single-cell rna-seq data. Briefings in Bioinformatics, 23(5):bbac313, 2022.

49. Junyuan Xie, Ross Girshick, and Ali Farhadi. Unsupervised deep embedding for clustering analysis. In International conference on machine learning, pages 478–487. PMLR, 2016.

50. Zehao Xiong, Jiawei Luo, Wanwan Shi, Ying Liu, Zhongyuan Xu, and Bo Wang. scgcl: an imputation method for scrna-seq data based on graph contrastive learning. Bioinformatics, 39(3):btad098, 2023.

51. Liying Yan, Mingyu Yang, Hongshan Guo, Lu Yang, Jun Wu, Rong Li, Ping Liu, Ying Lian, Xiaoying Zheng, Jie Yan, et al. Single-cell rna-seq profiling of human preimplantation embryos and embryonic stem cells. Nature structural & molecular biology, 20(9):1131–1139, 2013.

52. Fan Yang, Wenchuan Wang, Fang Wang, Yuan Fang, Duyu Tang, Junzhou Huang, Hui Lu, and Jianhua Yao. scbert as a large-scale pretrained deep language model for cell type annotation of single-cell rna-seq data. Nature Machine Intelligence, 4(10):852–866, 2022.

53. Bin Yu, Chen Chen, Ren Qi, Ruiqing Zheng, Patrick J Skillman-Lawrence, Xiaolin Wang, Anjun Ma, and Haiming Gu. scgmai: a gaussian mixture model for clustering single-cell rna-seq data based on deep autoencoder. Briefings in bioinformatics, 22(4):bbaa316, 2021.

54. Luke Zappia, Belinda Phipson, and Alicia Oshlack. Splatter: simulation of single-cell rna sequencing data. Genome biology, 18(1):174, 2017.

55. Amit Zeisel, Ana B Muñoz-Manchado, Simone Codeluppi, Peter Lönnerberg, Gioele La Manno, Anna Juréus, Sueli Marques, Hermany Munguba, Liqun He, Christer Betsholtz, et al. Cell types in the mouse cortex and hippocampus revealed by single-cell rna-seq. Science, 347(6226):1138–1142, 2015.

56. Zhenhao Zhang, Yuxi Liu, Jiang Bian, Antonio Jimeno Yepes, Jun Shen, Fuyi Li, Guodong Long, and Flora D Salim. Boosting patient representation learning via graph contrastive learning. In Joint European Conference on Machine Learning and Knowledge Discovery in Databases, pages 335–350. Springer, 2024.

57. Tong Zhao, Wei Jin, Yozen Liu, Yingheng Wang, Gang Liu, Stephan Günnemann, Neil Shah, and Meng Jiang. Graph data augmentation for graph machine learning: A survey. arXiv preprint arXiv:2202.08871, 2022.

58. Grace XY Zheng, Jessica M Terry, Phillip Belgrader, Paul Ryvkin, Zachary W Bent, Ryan Wilson, Solongo B Ziraldo, Tobias D Wheeler, Geoff P McDermott, Junjie Zhu, et al. Massively parallel digital transcriptional profiling of single cells. Nature communications, 8(1):14049, 2017.

59. Zilu Zhou, Bihui Xu, Andy Minn, and Nancy R Zhang. Dendro: genetic heterogeneity profiling and subclone detection by single-cell rna sequencing. Genome biology, 21:1–15, 2020.

